# Mapping the landscape of magnetic field effects on neural regeneration and repair: a systematic review, mathematical model, and meta-analysis

**DOI:** 10.1101/2023.03.14.532596

**Authors:** Meghan McGraw, Gabrielle Gilmer, Juliana Bergmann, Vishnu Seshan, Kai Wang, David Pekker, Michel Modo, Fabrisia Ambrosio

**Author notes:** These authors contributed equally and are considered co-first authors (Meghan McGraw, Gabrielle Gilmer). **Corresponding author:** Fabrisia Ambrosio, PhD, MPT, Office 5304, 149 13th St #4002, Charlestown, MA 02129.

## Abstract

Although magnetic field exposure is a well-established diagnostic tool, its use as a therapeutic in regenerative medicine is relatively new. Our goal here was to evaluate how magnetic fields affect neural repair *in vitro* by performing a systematic review of the literature, mathematical modeling, and meta-analyses. The 38 included articles presented with high heterogeneity, representing 13 cell types, magnitudes ranging from 0.0002-10,000 mT, frequencies from 0-150 Hz, and exposure times lasting from one hour to several weeks. Mathematical modeling revealed that increasing magnetic field magnitude increases neural progenitor cell (NPC) viability. For regenerative processes that were not influenced by magnitude, frequency, and time, we integrated data with meta-analyses. Results revealed that magnetic field exposure increases NPC proliferation while decreasing astrocytic differentiation. Collectively, our work identifies neural repair processes that may be most responsive to magnetic field exposure and provides a framework for novel hypothesis and technology development.

## Introduction

Magnetic fields are a ubiquitous part of life,^1^ exerting their effects and interactions at all scales, from subatomic particles to the universe itself.^2^ Organisms are continuously under the influence of exogenous magnetic fields (i.e., those produced by a sources outside of the body), which includes static fields such as Earth’s magnetic field^3^ as well as electromagnetic fields, such as those produced by modern-day power lines, electrical equipment, and cell phones.^4,5^ In addition, the human body itself produces magnetic fields, with fluctuations in cellular ion transport generating currents that ultimately create endogenous magnetic fields (i.e., those produced within tissues and organs themselves).^6^ One such organ in which magnetic fields are abundant is the brain.^7^ Neurons utilize sodium and potassium to generate action potentials for communication via the release of neurotransmitters. Modeling experiments have demonstrated that this ion movement likely leads to magnetic fields that may be experienced by neighboring cells (within 2-6μm).^8^

In recent years, considerable interest has turned to not only understanding endogenous magnetic fields but also toward understanding how exogeneous magnetic fields can be applied for therapeutic purposes.^9^ Magnetic field exposure as a treatment modality is appealing since it may be applied noninvasively and in the absence of anesthesia or ventilatory support. Indeed, transcranial magnetic stimulation (TMS) is already widely applied for the treatment of a variety of psychiatric disorders.^10^ The potential of applied magnetic fields has also been explored in the context of cerebral ischemic stroke, as highlighted by Moya-Gomez et al. in their review of electromagnetic field (EMF) applications in stroke from the culture dish to the clinic.^11^ Magnetic fields are thought to provide neuroprotective (i.e., rescue of apoptotic cells) effects,^11^ but the underlying mechanism remains unknown. Hypothesized mechanisms that may underlie physiological responses to magnetic fields include modulation of calcium and nitric oxide concentration,^12,13^ free radical production,^14^ inhibition of apoptosis,^15^ induced angiogenesis,^16^ manipulation of cellular electrical activity,^17^ and reduced edema/inflammation.^11,18^

Although magnetic field exposure appears to be beneficial in the context of stroke and some psychiatric disorders, there are other diseases where the impacts of magnetic field exposure are more ambiguous, such as Alzheimer’s disease. Some case-controlled and cohort studies report an increased risk of Alzheimer’s disease with static (no oscillations) magnetic field exposure (~10 uT).^19–21^ On the other hand, dynamic (oscillations) magnetic fields (10 mT, 100 Hz) increase brain activity in animal models of Alzheimer’s disease, in which it is hypothesized that protection results from the breakdown of β-amyloid plaques.^22^ Yet, just as in the case of stroke, the underlying mechanism of action remains unknown.

Several factors contribute to both the uncertainty in the benefit versus risk potential of magnetic fields as well as the lack of mechanistic insight. Magnetic fields have many tunable parameters, and the literature aimed at characterizing magnetic field effects on biological tissues is fragmented based on these parameters. Indeed, magnetic field properties (magnitude, frequency, pulsing), timing schedules (constant or fixed interval), species, system level (cell, tissue, organism), outcome of interest (proliferation, reproduction, cell health, organism behavior), and many other characteristics are highly variable across the literature. Given these characteristics undoubtedly dictate the therapeutic potential of magnetic fields, such study heterogeneity precludes drawing well-grounded conclusions. An improved understanding of the interactions between magnetic field variables and the corresponding mechanisms would aid scientists and clinicians in the systematic design of safe, predictable, and effective magnetic field therapeutics in humans.

Through a systematic review of the literature and application of meta-analytical tools, it is possible to improve our understanding of magnetic field effects on biological systems. Specifically, here, we performed a systematic literature search of studies pertaining to magnetic field effects on neural cell repair *in vitro*. We chose to focus on *in vitro* studies to expand on our mechanistic understanding of magnetic field effects on neural repair, as it is challenging to evaluate cell-based effects *in vivo* given the complexity of cellular interactions within the nervous system. We then compiled quantifiable repair characteristics (differentiation, proliferation, viability, maturation) as compared to controls, as well as magnetic field parameters (magnitude, frequency, timing of exposure) across included papers. With this information, we performed mathematical modeling and meta-analyses to characterize the effects of magnetic fields on neural repair. Additionally, we performed rigor and reproducibility analyses on the publications included in our review. Collectively, our work identifies magnetic field parameters that may be most effective in eliciting pro-regenerative responses, as well as cellular repair processes that are responsive to any sort of magnetic stimulation.

## Methods

### Systematic Review

We conducted a systematic search of the literature using the Preferred Reporting Items for Systematic Reviews and Meta-Analyses (PRISMA) statement and protocol (**Appendix S1**),^23,24^ the Meta-Analysis of Observational Studies in Epidemiology (MOOSE) checklist (**Appendix S2**),^25^ the Cochrane Handbook for Systematic Reviews of Interventions,^26^ and the practical guide for meta-analysis in animal studies.^27^ We set our PECO (<u>P</u>opulation, <u>E</u>xposure, <u>C</u>omparison, <u>O</u>utcome) inclusion criteria to identify publications that studied human or mammalian neural cells *in vitro* (population) that were exposed to magnetic fields (exposure). In these publications, control conditions consisted of a matched neural cell type that was not exposed to magnetic fields beyond Earth’s magnetic field (comparison), while the outcome variable of interest was a metric of cellular repair or regenerative processes (outcome). Included studies had to be published in a peer-reviewed scientific journal and written in English. Publications were excluded if they studied the whole organism, other organismal tissues (e.g., bacteria, plant), non-neural cell types, or a high static magnetic field (as experienced in a magnetic resonance imaging (MRI) scanner).

We performed electronic searches of PubMed, Association for Computing Machinery (ACM) Library, Association for Information Systems eLibrary (AISeL), Institute of Electrical and Electronics Engineers (IEEE) Library, Springer, Web of Science, Scopus, Science Direct, and arXiv on September 15, 2021, with search terms outlined in **Table S1**. A manual search on Google Scholar was also performed. Afterwards, we performed title and abstract screening in ASReview, which is a machine learning algorithm for time-efficient and reproducible screening of titles and abstracts.^28^ Full-text screening was performed by two independent reviewers (MM, JB) based on the Cochran Handbook recommendations.^26^ Afterwards, we performed a citation search of all included publications using Web of Science.^26^ All screening steps were completed in pre-designed spreadsheets, and disagreements between reviewers were adjudicated by a third reviewer (GG).

One reviewer (MM) extracted study information from the included publications (title, cell type, outcome variable category, specific outcome variable, sample size, magnitude, frequency, total exposure time, time after exposure). When referring to the different cell types, throughout this manuscript we use the term ‘neural’ to mean any cell that makes up the nervous system, ‘neuronal’ to refer to cellular functions related to neurons, and neural stem/progenitor cell (NPC) to refer to neural stem cells, induced pluripotent stem cells if they were maintained as a stem cell in the experiment, and neural progenitor cells. Specific outcome variables were then categorized into the following regenerative processes: viability, proliferation, differentiation, and maturation. ‘Viability’ was defined as a metric that measured the ability of the cell to survive. ‘Proliferation’ was defined as a metric that measured increases in the number of cells or signs of an active cell cycle (i.e., mitosis). ‘Differentiation’ was defined as a metric that tracks the transformation from stem/progenitor cell to a specific mature cell type. ‘Maturation’ was defined as a metric that quantified cellular size. Some dependent variables were included under multiple categories if that metric was used to categorize different repair processes. For example, Glial Cell Line-Derived Neurotrophic Factor (GDNF) is known to be associated with increased viability^29^ and differentiation,^30^ and it was therefore included in both sets of mathematical models. Graphical data were converted to numerical data via a digital ruler. Intra-rater reliability for graphical to numerical data conversion, calculated as the reviewer measuring the same data in sessions ten months apart (n = 5), was determined to be ‘excellent’ (intraclass correlation coefficient [2,1]: 0.9999 95% CI: [ 0.9846, 1.000]).

### Rigor and Reproducibility Assessment

We scored the rigor and reproducibility of the included manuscripts using the ARRIVE guidelines 2.0.^31^ Two independent reviewers (MM and GG) ranked the categories comprising the ARRIVE guidelines as either ‘clearly insufficient’ (0), ‘unclear if sufficient’ (1) or ‘clearly sufficient’ (2). Any disagreements in ranking that differed by 2 were discussed until a consensus was reached. Otherwise, the results between the two reviewers were averaged. Since our analysis was limited to *in vitro* studies, conditions 14-16 (“Ethical Statement”, “Housing and Husbandry”, “Animal Care and Monitoring”) were not included in this assessment. The maximum score was a 36, indicating higher rigor, whereas the lowest score possible was a 0, indicating a complete lack of rigor. Individual categories (e.g., “Randomization”) were considered ‘sufficiently reported’ if they had an average score > 1 and a standard deviation < 0.25, whereas ‘insufficiently reported’ was equated to an average score < 1 and a standard deviation < 0.25. An ‘unclear if sufficiently reported’ was recorded if the category score did not fall into these two categories. To test our hypothesis that articles published more recently would have higher ARRIVE scores, SPSS Statistics for Windows (version 28.0, IBM Corp., Armonk, NY, USA) was used to analyze the correlation between a study’s ARRIVE score and its year of publication.

### Mathematical Models

The included studies offered varying ranges of properties and exposure schedules of magnetic fields as well as a variety of metrics to quantify repair and regenerative processes. In categorical (standard) meta-analysis subjects can be divided into two groups: experiment and control. However, the exposure schedules in the dataset we analyzed were not categorical but, continuously-varying. Therefore, for the first pass through the dataset, we performed a series of mathematical models, similar in structure to continuous exposure meta-analyses,^32^ in order to (1) extract the continuous relationship between exposure and regenerative processes and (2) derive information that cannot be extracted from individual studies. We tested a variety of mathematical models to evaluate how different magnitudes, frequencies, percent exposure times, and total exposure times (magnetic field parameters, independent variables) affected viability, proliferation, differentiation, and maturation of neural cell populations (repair and regenerative process, dependent variables). Magnitude is defined as the strength or amplitude of the magnetic field, and frequency is defined as the number of cycles of magnetic field per unit time. The percent exposure times represents the proportion of experiment time that cells were exposed to a magnetic field relative to the total time of the experiment. For example, if cells were exposed to a magnetic field for 1 hour, but 10 hours passed from the start of the experiment to data collection, the percent exposure time would be 0.1. The total exposure time is the amount of time exposed to the magnetic field (in the previous example, the total exposure time is one hour).

We note that none of the studies controlled for the effects of Earth’s static magnetic field (~50 μT). Instead, the control group in all included studies consisted of unmodified exposure to Earth’s magnetic field. On the other hand, the experimental group consisted of Earth’s static magnetic field plus an additional applied field. As all of the studies reported magnetic field amplitude but not magnetic field vector data, we did not have sufficient information available to compensate for the Earth’s magnetic field (because magnetic fields add as vectors and not scalars). Therefore, we ignored the contribution of the Earth’s magnetic field in our models. Specifically, we set the magnetic field to zero for the control group and did not compensate the exposure groups for Earth’s magnetic field. The lack of vectorial magnetic field data makes our mathematical model insensitive to magnetic field effect that occur when magnetic field amplitudes are on the order of Earth’s magnetic field or lower.

The mathematical model described below was run only if there were at least 6 different values of the independent variable (e.g., magnitudes of 0.1, 1, 2, 5,10, and 100 mT versus all datapoints investigating 100 mT). All simulations were completed in Python Jupyter Notebook (version 6.3.0) and followed recommendations for best practices in mathematical modeling.^33,34^ For each independent– dependent variable combination, initial settings for the mathematical model (linear and polynomial) and initial coefficient estimates were generated via the *numpy polyfit* function (Python 1.23.0).

To set up the simulations, the ideal model was set up as follows:

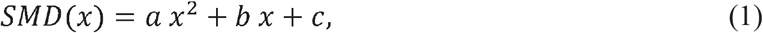

where *x* is one of the independent variables (e.g., magnetic field magnitude); *SMD(x)* is the standardized mean difference between the magnetic field exposure group and the control group from an individual study for a repair/regenerative process dependent variable category; *a* and *b* are the coefficients predicted by the simulation; and *c* was set to 0 (i.e., SMD = 0 for the control group). Therefore, an SMD of zero would indicate there is no difference between the magnetic field exposure and control group, a positive SMD would indicate that the specific dependent variable is greater in the magnetic field exposure group compared to the control group, and a negative SMD would indicate that a specific dependent variable is greater in the control group compared to the magnetic field exposure group. For specific variables that are inversely correlated with their category, the sign of the SMD value was reversed. For example, while BrdU is positively correlated with proliferation,^35^ p53 is inversely correlated with proliferation.^36^ In the case of BrdU, an increase in the experimental group compared to the control would yield a positive SMD, while for p53, this would result in a negative SMD. As such, the sign of the p53 SMD would be flipped from negative to positive for the purpose of compiling all metrics together. This nuance prevents the following situation: if there is a variable that is known to be 1:1 positively correlated with the independent variable (y = x), while another variable that is known to be 1:1 negatively correlated with the independent variable (y = -x), upon combination, there will be no correlation (y = 0). By reversing the negatively correlated function, we can simply track changes as related to the expected behavior of the variable, with positive being in the direction of enhanced activity, and negative being in the direction of attenuated activity.

In our analysis, we fitted the coefficients of Eq (1) by minimizing the root mean squared error (RMSE). For articles that included multiple exposure settings, we included each condition from that dataset as an independent input into our model. To accommodate the fact that some exposure settings appeared in multiple different studies we ran 20,000 different fits. In each fit, if inputs were the same across different studies, one was randomly selected. For instance, if study A used a magnitude of 10 mT and study B also used a magnitude of 10 mT, whether the SMD for study A or study B was input into the model was randomly selected. To ensure that we were adequately sampling identical inputs, we tracked how the conclusions drawn from the analysis varied with the number of separate fits. We observed that 200 separate fits were sufficient to fix the values of a and b. Finally, as stated above, we assumed that an external magnetic field exposure of zero (magnitude, frequency, time) would be equivalent to the control and input this value as an SMD of 0.

The null hypotheses for our simulations were that there is no relationship between our independent variables (magnitude, frequency, exposure time) and repair processes (viability, proliferation, differentiation, maturation). In our models, this would present as coefficients (i.e., *a, b)* equal to zero. To evaluate this statistically, we defined alpha *a priori* as 0.05. The p-values of our simulations were quantified as the number of values greater than or less than zero divided by the total number of completed simulations. For example, if the average value of *b* was 9.82 and three out of 100 completed fits produced a negative value of *b*, our p-value would be 0.03 and this finding would be considered statistically significant.

After we ran the fits, we considered which datasets had significant *a* and/or *b* values and completed a series of sensitivity analyses to test the robustness of these findings. We first perturbed the *a* and/or *b* values from 10 – 200% of their original values and reconverged the RMSE to quantify how the model was affected by changes in *a* and *b*. If the RMSE was affected by changes in both *a* and *b* or just *a*, then the polynomial model was analyzed. If just *b* affected the RMSE, only the linear model was analyzed. We then altered *a* and/or *b* values again (increased 5 – 20%) to observe the effects on predicted SMD and to analyze the robustness of our model as a function of error in our coefficients. We marked a model as robust if a 20% change in *a* and/or *b* resulted in less than an absolute value change of 2 for the SMD. The original codes for all simulations are shown in **Appendix S3**.

### Categorical meta-analyses

The continuous exposure meta-analysis model offers the advantage of identifying the continuous relation between dependent and independent variables. However, it is not as sensitive as categorical meta-analysis which is designed to test a binary hypothesis.^32^ Therefore, we performed categorical meta-analysis for the combinations of dependent (e.g. oligodendrocytic differentiation, astrocytic differentiation, etc.) and independent variables (magnetic field amplitude, frequency, and exposure time) for which we did not find a continuous relation. In each case the control group was chosen to have no external magnetic field exposure and the experimental group was chosen to include all cases with external magnetic field exposure. SMD and pooled standard deviations (SD_pool_) of outcome measures were calculated using the DerSimonian-Laird method.^26^ All meta-analyses were performed using SPSS Statistics for Windows (version 29.0, IBM Corp., Armonk, NY, USA).

## Results

### Systematic review revealed a wide variety in magnetic field parameters, cell types, and outcome measures for neural repair processes

From our systematic review, we identified 7,917 articles that evaluated the effects of magnetic field exposure on repair processes in neural cells *in vitro*. Title and abstract screening via ASReview, which is a machine learning algorithm for large review screening, resulted in 159 articles eligible for full-text screening **(Table S2).** After full-text screening and a citation search, 38 articles were included in this review (**Figure 1**, **Table S3**).^37–73^ The most frequent reason for exclusion was whole animal exposure to magnetic fields (**Table S2**).

**Figure 1:**
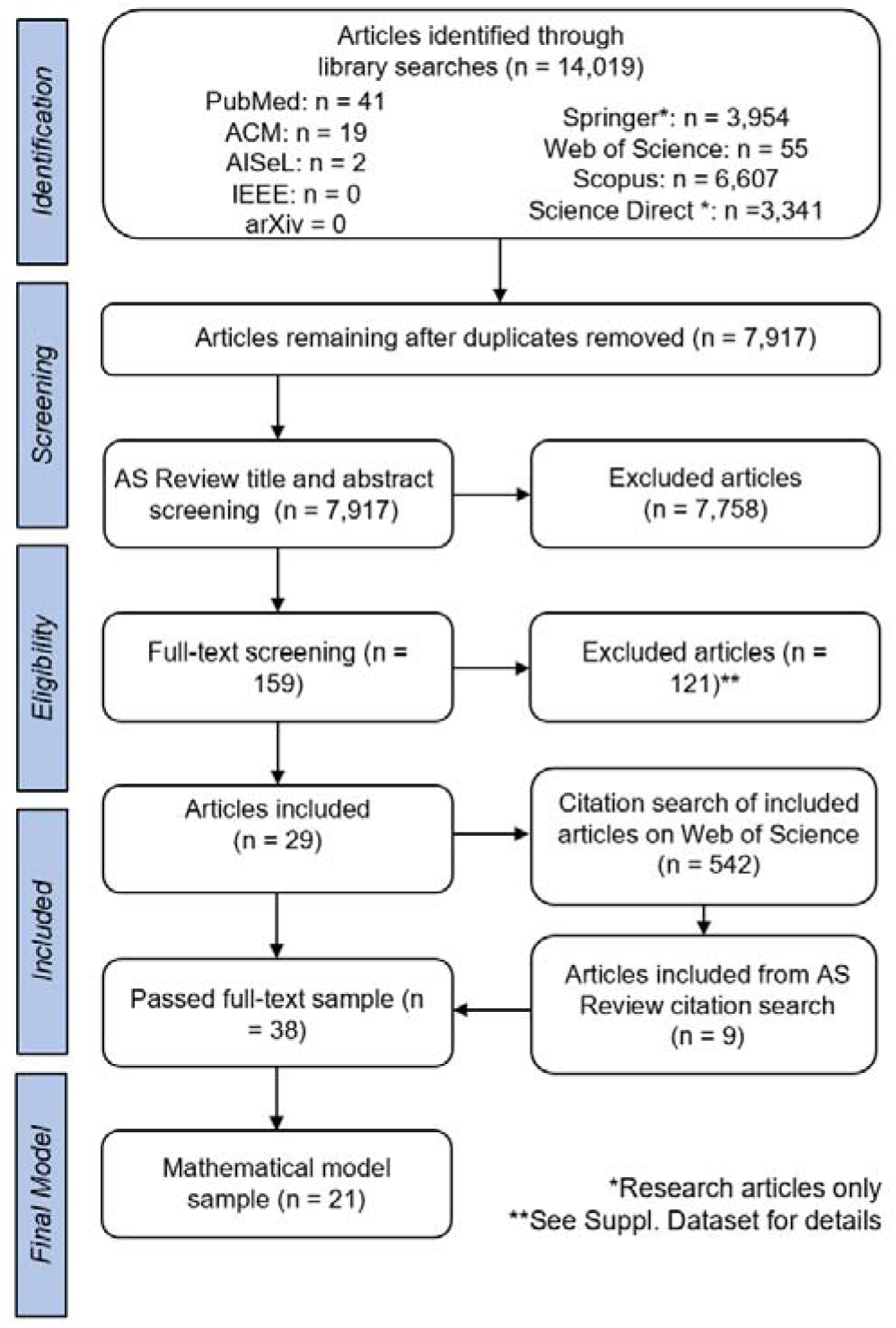
Systematic review workflow. Systematic search of databases, abstract, title and full text screening as well as a citation search resulted in 38 unique articles included in this review. Details on exclusion reasons at the full text level and included article details are available in Tables S2 and S3.

Of the studies that were included, 35 used murine and 4 used human cells (1 study used both; **Figure 2A**). **Figure 2B** displays the breakdown of cell types used in each article, with neural progenitor cells (n = 11) being the most common. When considering magnetic field parameters, dynamic (n = 21, **Figure 2C**), low magnitude (1 - 10 mT, n = 30, **Figure 2D**), with a 50 Hz frequency (n = 17, **Figure 2E**) fields were the most frequently used. When considering outcome variables, most studies examined how magnetic field exposure affected differentiation and proliferation (**Figure 2H**). The total time of exposure to the magnetic field was variable, ranging from less than one hour to over one week (**Figure 2F**). The total time of exposure compared to the total length of the experiment demonstrated a bimodal distribution, with one cluster of studies focusing on exposure < 10% of the total experiment length (e.g., one quick burst at the beginning of the experiment) and the other cluster of studies exposing cells for nearly the entire length of the experiment (>90%) (**Figure 2G**). These findings reveal a high amount of heterogeneity in studies aimed at understanding how magnetic field exposure impacts neural cellular repair processes.

**Figure 2:**
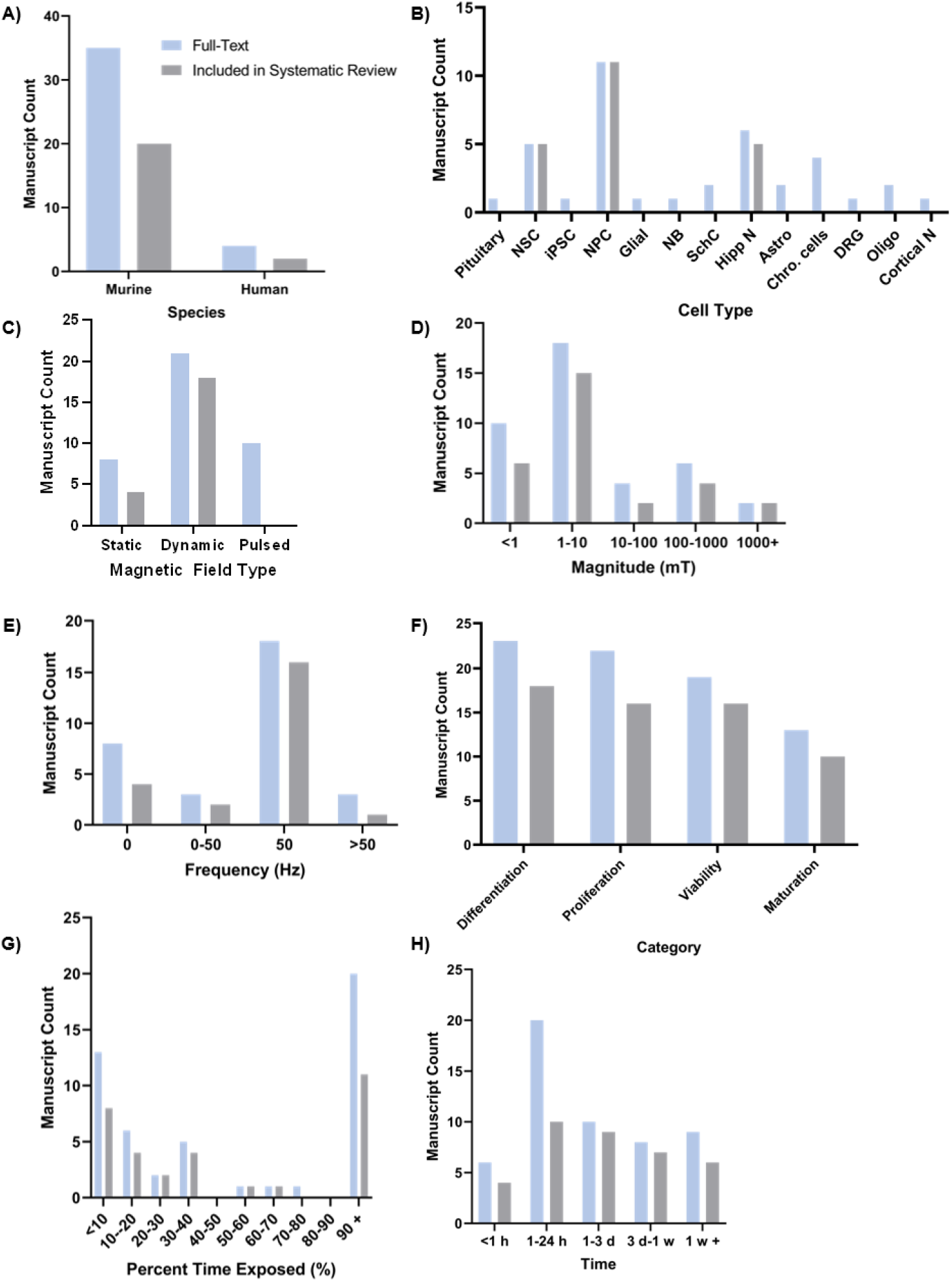
Publications aimed at studying neural repair as a function of magnetic field exposure offer a wide range of diversity in both magnetic field parameters and cell types under study. A) Species of cells. B) Cell Type. NSC = Neural Stem Cells. iPSC = Induced Pluripotent Stem Cells. NPC = Neural Progenitor Cells. NB = Neuroblastoma. SchC = Schwann Cells. Hipp N = Hippocampal Neurons. Astro = Astrocytes. Chro Cells = Chromatin Cells. DRG = Dorsal Root Ganglia. Oligo = Oligodendrocytes. Cortical N = Cortical Neurons. C) Type of magnetic field. D) Magnitude of magnetic field. E) Frequency of magnetic field. F) Metric of repair/regeneration studied. G) Percent of time exposed (time exposed to magnetic field over total time of experiment and outcome measurement). H) Total time of magnetic field exposure.

### Included studies consistently lacked information on “Inclusion and Exclusion Criteria”, “Protocol Registration”, and “Data Access”

To assess rigor and reproducibility (R&R), we scored each manuscript by ranking the categories within the ARRIVE guidelines. Included manuscripts had an average R&R of 23±4, (**Figure 3, Table S4** includes scoring per manuscript). The maximum overall score, indicating higher rigor, is 36, and the lowest score, indicating no rigor, is zero (see **Methods** for details). When examining the Essential 10, “Study Design”, “Outcome Measures”, “Experimental Procedures”, and “Results” were ‘sufficiently reported’. On the other hand, “Inclusion and Exclusion Criteria” was ‘insufficiently reported’. Out of the additional guidelines, “Abstract”, “Background”, and “Objectives” were ‘sufficiently reported’, while “Protocol Registration” and “Data Access” were scored as ‘insufficiently reported’ across all studies (**Figure 3A**). To determine if the level of rigor and reproducibility improved over time, we evaluated the correlation between the ARRIVE Score and year of publication (**Figure 3B**). We found there was a moderate, positive correlation, suggesting that R&R is slowly but steadily improving over time.

**Figure 3:**
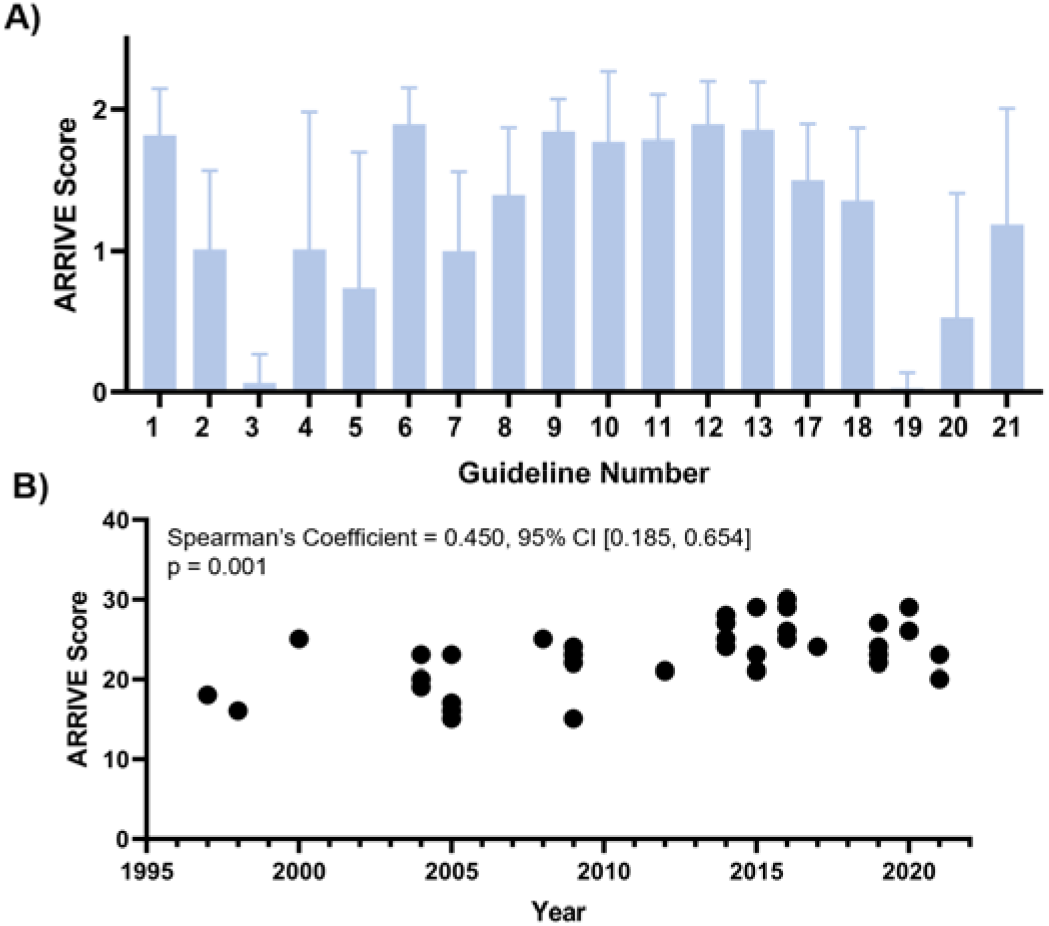
Rigor and reproducibility in studies aimed at understanding magnetic field impacts on neural repair is increasing across time. A) ARRIVE score based on individual category, with 2 being sufficiently addressed and 0 being insufficiently addressed. 1 = Study Design. 2 = Sample Size. 3 = Inclusion & Exclusion Criteria. 4 = Randomization. 5 = Blinding. 6 = Outcome measures. 7 = Statistical methods. 8 = Experimental animals. 9 = Experimental procedures. 10 = Results. 11 = Abstract. 12 = Background. 13= Objectives. 17 = Interpretation/scientific implications. 18 = Generalizability / translation. 19 = Protocol registration. 20 = Data access. 21 = Declaration of interests. B) Relationship between total ARRIVE Score and publication year.

When looking qualitatively at factors driving our R&R results, we found that most studies listed the sample size but did not report power analyses that ensured sample size was appropriate for the statistical analyses (n = 33; listed as number of studies that did not comply). We also found that some, but not all, procedures were randomized (n =19) and blinded (n = 15). While almost every study reported the statistical tests performed, few cited whether their datasets met the appropriate assumptions (n = 29). Some studies did not clearly articulate the limitations of their studies (n=27) or the clinical translatability (n = 27), no study registered their protocols (n = 38), and most studies did not report a means for accessing the data (n = 27). With regards to declaration of interests, many studies did not include the grants under which they were funded (**n** = 18). Lastly, no study included descriptions of the passage number of cells, or the age or sex of the animal from which cells were derived (n = 38).

### Mathematical modeling reveals magnetic field magnitude has a stable, positive relationship with neural stem/progenitor cell (NPC) viability

To begin interrogating how magnetic field exposure modulates neural repair, we performed a series of mathematical models simulating the effects of magnetic field parameters (magnitude, frequency, total time of exposure, percentage of time exposed) on metrics of neural repair (viability, proliferation, differentiation, maturation). Studies that did not contain outcome measures falling into one of these categories or studies that used a pulsed magnetic field paradigm were excluded to simplify the model input. Twenty-one articles met these guidelines and were included in subsequent analyses (study characteristics outlined in **Figure 2;** the modeling workflow is displayed in **Figure 4A**; the model input is presented in **Table S5**).

**Figure 4:**
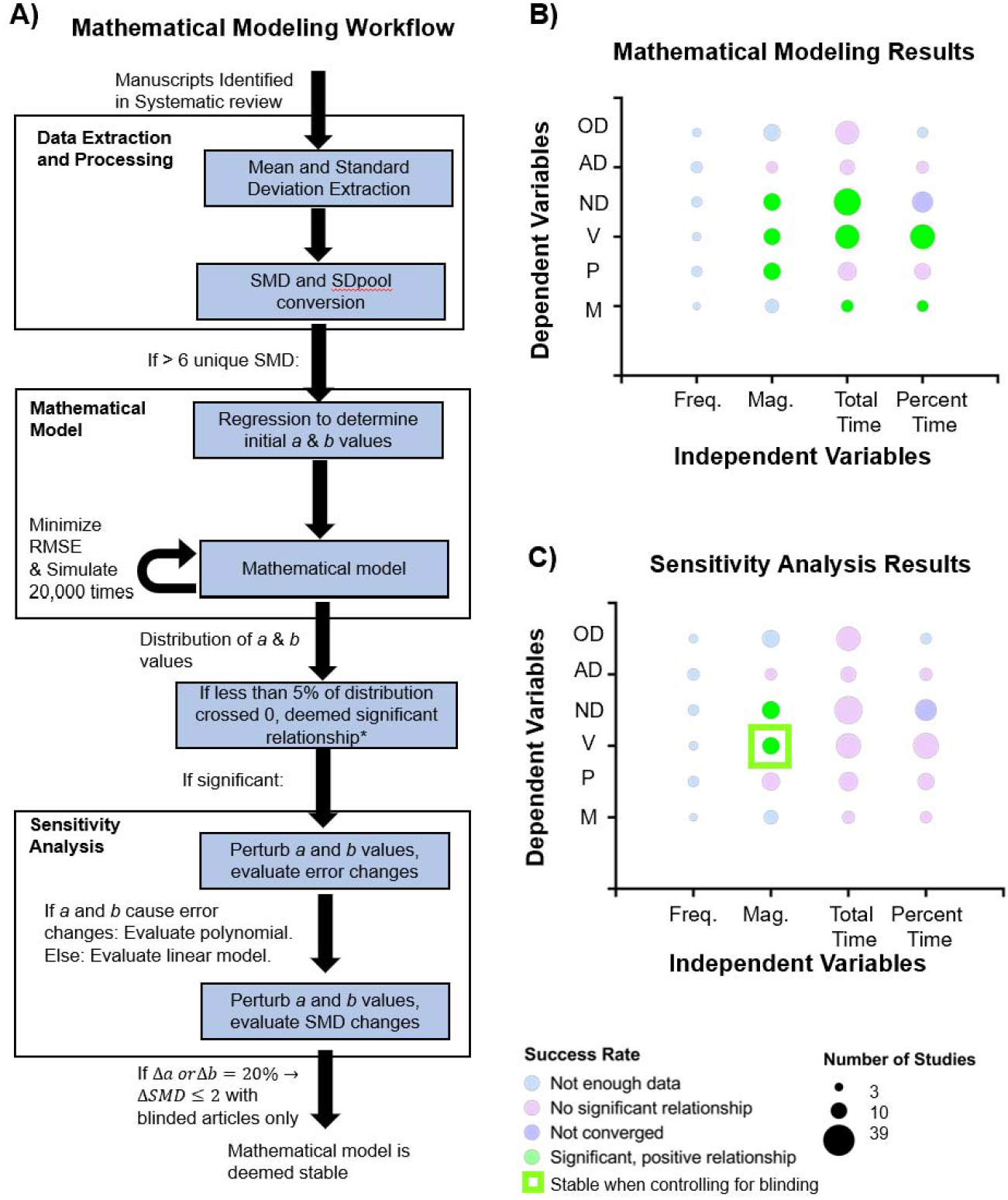
Mathematical modeling and sensitivity analyses reveal NPC viability has a robust relationship with magnitude of magnetic field exposure. A) Mathematical modeling and sensitivity analyses workflow. SMD = Standardized Mean Difference. RMSE = Root Mean Squared Error. SDpool = Pooled standard deviation. *For example, if the average value of *a* was 3 and only 3% of simulated *a* values were negative, we would say with confidence that there is a relationship between the independent and dependent variables. B) Results from mathematical modeling. OD = Oligodendrocytic differentiation. AD = Astrocytic differentiation. ND = Neuronal differentiation. V = Neural Stem/Progenitor Cell (NPC) Viability. P = NPC Proliferation. M = NPC Maturation. Mag. = Magnitude. Freq. = Frequency. Of note, no negative relationships were observed in our modeling results. C) Sensitivity analysis results.

After quantification, six distinct dependent variables were tested: neuronal, astrocytic, and oligodendrocytic differentiation, as well as neural stem/progenitor cell (NPC) maturation, proliferation, and viability. Considering the independent variables (magnitude, frequency, total time, percent time), a total of 24 combinations were possible for simulation. However, nine combinations did not have enough data (a minimum of 6 unique levels of independent variable inputs) to run a simulation. Thus, 15 models were simulated (**Figure 4A, Table S6-S7**).

Based on our models, astrocytic differentiation, and oligodendrocytic differentiation did not have a relationship with magnitude, frequency, time, or percentage of time exposed to the magnetic field **(Figure 4B**). Changes in time-based variables were positively correlated with neural maturation, while NPC proliferation was positively correlated with increasing magnitude. NPC viability had significant, positive correlations with respect to magnitude, total time, and percent time. Lastly, neuronal differentiation was positively correlated with magnitude and total time exposed. Taken together, these simulations suggest that magnetic field exposure may exert positive effects on NPC maturation, proliferation, differentiation, and viability.

To assess the robustness of our simulations, we performed a series of sensitivity analyses. First, we analyzed how changes in *a* and *b* values affected the RMSE. For most cellular repair processes, we found that both *a* and *b* affected the RMSE. Therefore, we modeled polynomial relationships for the entirety of the sensitivity analyses. For all considered combinations, RMSE effects were stable.

Unlike the statistically significant relationships observed in fitting to mathematical models, sensitivity analyses revealed a lack of robustness across many of the observed relationships, as quantified by a change in SMD > 2 (**Figure 4C**). Models assessing the relationship between NPC viability and neuronal differentiation with respect to magnitude were stable. The total time models generally showed moderate instability (SMD of 2-10 for both *a* and *b*), while the percent time models showed extreme instability (SMD of ~10^5^ for *a* and ~10^3^ for *b*).

Using the previous data generated from our rigor and reproducibility analysis, we repeated the mathematical models only including those that explicitly used blinding in their experiments (in other words, a score of 2 in the 5^th^ ARRIVE Guideline Category, ‘Blinding’). We found that 6 manuscripts used in the mathematical models met this criterion. Of note, astrocytic differentiation and NPC maturation simulations could not be run because there were no independent variables with at least 6 unique datapoints. Neuronal differentiation, oligodendrocytic differentiation, and NPC proliferation only had enough independent variables for total time, while NPC viability had enough data for both total time and magnitude simulations. Oligodendrocytic differentiation and NPC proliferation did not have a relationship with time exposed to the magnetic field. Changes in total time were positively correlated with neuronal differentiation and NPC viability, while changes in magnitude were positively correlated with NPC viability. However, the neuronal differentiation model and total time model did not converge. Sensitivity analyses revealed that the relationships between NPC viability with respect to total time and magnitude were stable. Combining this with our original mathematical modeling results, our findings indicate with confidence that there is a positive relationship between magnetic field magnitude and NPC viability (**Figure 4C**; raw data shown in **Figure S1**).

### Meta-analyses reveal that NPC proliferation is increased, whereas astrocytic differentiation is decreased by magnetic field exposure

Based on our mathematical modeling results, oligodendrocytic and astrocytic differentiation, as well as NPC proliferation were not modulated by changes in magnitude, frequency, or time of exposure to a magnetic field. Because there were no apparent effects of these heterogeneous exposure parameters, we next compiled these datasets to further characterize whether there were any effects of magnetic field exposure versus no exposure via meta-analysis.

Categorical meta-analysis revealed that NPC proliferation was increased in samples exposed to magnetic fields compared to samples that were not exposed (**Figure 5A**). Conversely, astrocytic differentiation was decreased (**Figure 5B**). Lastly, oligodendrocytic differentiation was not impacted, although this analysis is likely underpowered, as there were only three independent studies included (**Figure 5C**). Collectively, these findings suggest that magnetic field exposure may amplify some aspects of neural repair/regeneration, such as NPC proliferation, while dampening other aspects, such as astrocyte differentiation.

**Figure 5:**
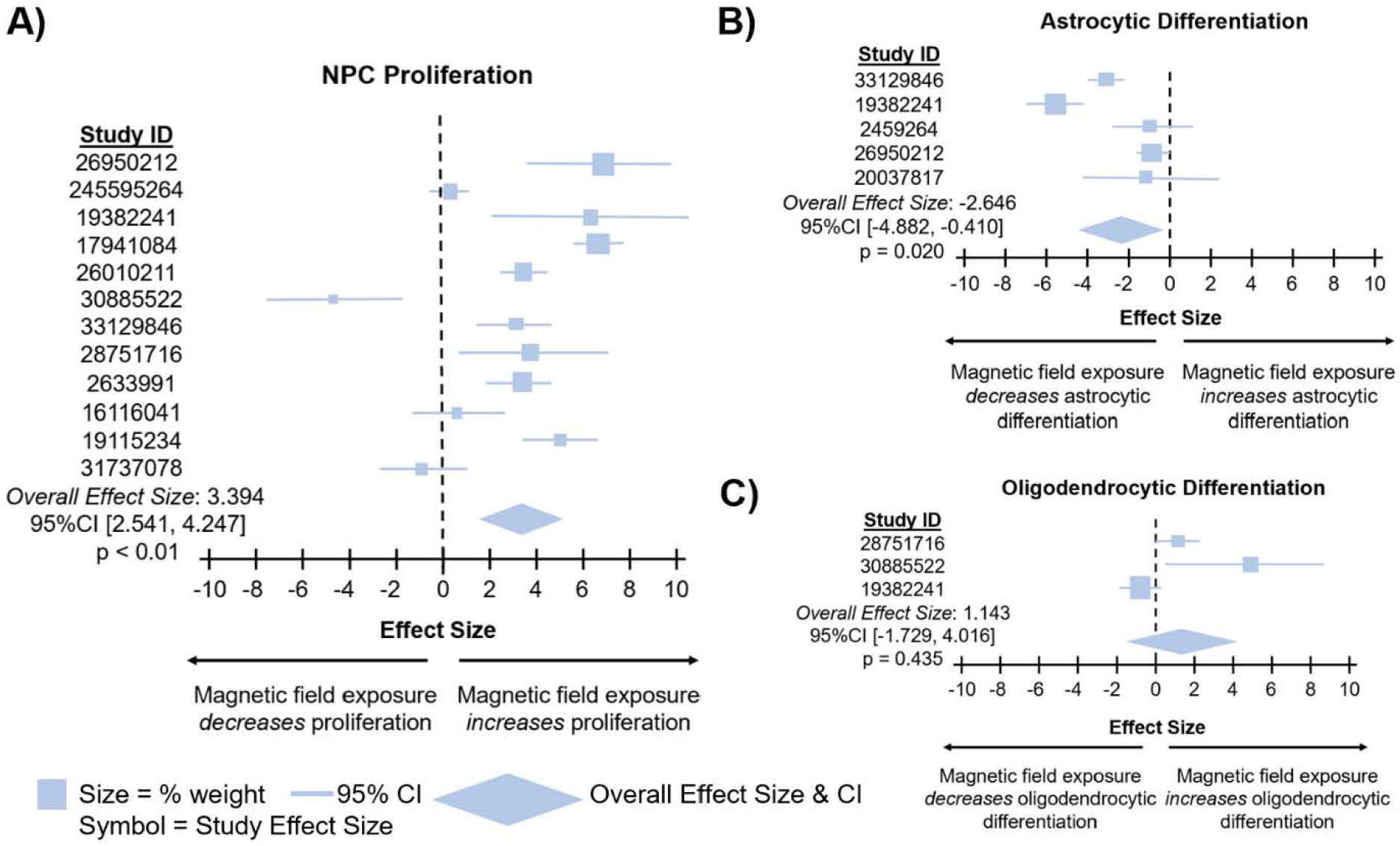
Meta-analyses revealed that magnetic field exposure increases NPC proliferation and decreases astrocytic differentiation. A) Meta-analysis assessing neural stem/progenitor cell (NPC) proliferation from included studies. B) Meta-analysis assessing astrocytic differentiation from included studies. C) Meta-analysis assessing oligodendrocytic differentiation.

## Discussion

As a noninvasive approach, magnetic field simulation represents a promising therapeutic in regenerative medicine, particularly for neural regeneration. To improve our understanding of magnetic field impacts on neural repair and regenerative cellular processes, we first cataloged all studies aimed at understanding the impact of magnetic field exposure on neural repair *in vitro* through a comprehensive literature search. From the identified studies, we then performed a series of mathematical models to evaluate how varying magnetic field parameters impacted neural repair. We further performed meta-analyses to characterize the effects of magnetic fields on neural repair. Our mathematical models revealed that increasing magnetic field magnitude increases NPC viability, while our meta-analyses revealed that general magnetic field stimulation increases NPC proliferation but decreases astrocytic differentiation.

In considering the magnetic field parameters tested (**Figures 2D-2E**), the studies aimed to simulate everyday exposures. For example, 50 Hz fields are reflective of those emitted by cell phones^74^ and power lines,^75^ and there are some human studies suggesting that this exposure has detrimental tissue effects.^76^ Other features, such as time of exposure, lacked a clear rationale. For example, when we sought to understand the wide range of exposure times investigated (**Figure 2G-H**), we found that authors infrequently reported why exposure schedules were chosen. Future work would benefit from a more clearly presented exposure schedule rationale and/or systematic evaluation of how magnetic field parameters impact repair/regenerative outcomes of interest.

In 2010, the first iteration of the ARRIVE Guidelines were published to provide a framework for scientists to address the rigor and reproducibility crisis that has become epidemic in the biological sciences.^77^ Clearly, progress is being made as evidenced by the positive correlation seen between publication year and ARRIVE Score (**Figure 3B**). However, in the body of literature aimed at understanding magnetic field effects on neural repair and regeneration, there are still many areas for improvement, including declaration of inclusion & exclusion criteria and registering protocols *a priori*. Although these shortcomings are not unique to this body of literature, given the high heterogeneity of study parameters (i.e., magnetic field specifications) and associated gap in understanding of ideal exposure conditions, increased attention to these features is essential.

While meta-analysis is the standard for evaluating overlapping parameters in systematic reviews, here, we applied mathematical modeling to better incorporate the heterogeneity present within the included studies. Our mathematical modeling results revealed that NPC viability improved with increasing magnetic field magnitude. This finding is consistent with several independent studies ^38,54,78^ and cell types. ^79–81^ Since neurons themselves lack a robust ability to regenerate,^82^ maintaining viability of NPC to restore support cells is of critical importance to neural recovery and repair. Our findings suggest that one mechanism of therapeutic benefit provided by magnetic fields is the protection of NPC. There is some evidence suggesting these effects may be modulated by interactions between the magnetic field and voltage-gated ion channels, leading to stabilization of cell membranes.^83,84^ Others suggest this improved viability may be modulated by magnetoelectric materials present within the cells that facilitate temperature control.^85,86^ Our findings here suggest that examination of these mechanisms to both optimize the therapeutic potential and better understand how magnetic field stimulation affects NPC viability is a promising direction for future research.

Our mathematical modeling suggested there were no effects of magnetic field magnitude, frequency, or exposure time on oligodendrocytic differentiation, astrocytic differentiation, or NPC proliferation. Thus, we evaluated the correlations of these magnetic stimulation parameters and cell features by performing categorical meta-analyses. Our meta-analyses indicated that magnetic field exposure may increase NPC proliferation, while decreasing astrocytic differentiation. Astrocytes play a critical role in neural repair/regeneration by activating NPC and creating an environment that allows for neuronal maturation and proliferation.^87^ However, overactive astrocytes can lead to glial scarring and serve as a detriment to repair.^88^ In other work exploring how magnetic fields affect astrocytes, authors show maturation phase affects how astrocytes respond, with exposure performed during differentiation enhancing astrocyte abundance.^89^ The differential response among varying cellular repair processes to magnetic field exposure may explain some of the inconsistencies in benefit versus risk that have been reported in human and animal studies. Additionally, there may be a ‘goldilocks effect’ in which some ideal magnetic field exposure increases NPC viability and proliferation, while preserving the appropriate proportion of astrocytes.^90^ As such, future studies are needed to systematically evaluate how these three parameters of magnetic field exposure—magnitude, frequency, and time—modulate regenerative/repair cascades.

We note that a lack of compensation for Earth’s magnetic field and the lack of vectorial magnetic field data make our analysis insensitive to effects of small magnetic fields (i.e., magnetic fields that have amplitudes similar or less than the Earth’s magnetic field). We hypothesize that biological systems that evolved in ~ 50 μT Earth magnetic field would be sensitive to magnetic field in the 1-100 μT range. In order to test this hypothesis it is important for the next generation of experiments to carefully compensate for the Earth’s magnetic field. As the magnetic field is a vector, compensating the Earth magnetic field requires a vector magnet (e.g., using three Helmholtz coils) and some way to measure the magnetic field in the x, y, and z direction (e.g., using a flux gate).

Although our analyses here add to a growing body of literature assessing the therapeutic potential of magnetic fields, they have limitations. First, the included studies were all performed *in vitro.* In addition, all of the studies were monocultures. Thus, there are likely a multitude of systemic interactions, such as the impact of microglial inflammation, not captured. In addition, 3D cultures tend to behave more analogously to *in vivo* systems than 2D cultures; however, all the studies in this review used 2D modeling. There were a diversity of cell types, species, outcome measures, and exposures included in this review, which impacts the results in a way that we are unable to account for. More specifically, the time and schedule of exposure were highly variable among studies and, thus, challenging to simplify for modeling purposes. Additionally, interest in pulsed electromagnetic fields is increasing due to its production by cell phones, but due to the quantification of the magnetic fields in this study, it was impossible to incorporate these more complex waveforms.^91^ It is also possible that there were impacts of recency here that were not fully elucidated (i.e., 1 hour exposure then sample collection 24 hours later versus 1 hour exposure and immediate sample collection). Given this variability, our models cannot be taken as an exact representation of existing relationships, but rather, a general, binary trend of whether a relationship exists or not. Most simulations and meta-analyses contained a relatively low number of studies. It is therefore possible some of our analyses were underpowered. The original data input in the mathematical models were usually clustered around one or a few points (e.g., magnitude tended to be close to 0 mT, while frequency was centered around 0 and 50 Hz). This observation stresses the need for more studies to evaluate uncommon parameters to better inform the therapeutic window of magnetic fields. Given these considerations, our analyses provide guidance to improve the design and reporting of magnetic field studies in the context of neural cell repair and regeneration. In addition, further improvement of our understanding of magnetic field parameters impact on cellular processes can lead to enhancing endogenous tissue repair and therapeutic interventions.

Magnetic field exposure will continue to be a part of everyday life and the potential for a noninvasive therapeutic will likely increase with our understanding of magnetic field effects on repair and regenerative processes. Future steps should include increased emphasis on systematic evaluation of magnetic field exposure that impacts NPC viability and proliferation and astrocytic differentiation in original research studies as well as the underlying mechanisms dictating these phenotypes. By increasing our understanding of the nuance of magnetic field effects on neural regeneration, we will move closer towards a robust and safe therapy.

## Author contributions

All authors made substantial contributions in the following areas: (1) conception and design of the study, acquisition of data, analysis and interpretation of data, drafting of the article; (2) final approval of the article version to be submitted; and (3) agreement to be personally accountable for the author’s own contributions and to ensure that questions related to the accuracy are appropriately investigated, resolved, and the resolution documented in the literature.

The specific author contributions are as follows:

GG, DP, and FA provided the concept, idea, and experimental design for the studies.

MM*, GG, and FA wrote the manuscript.

MM*, GG, VS, JB, and FA provided data collection, analyses, and interpretation of the data. MM*, GG, VS, JB, KW, DP, MM, and FA reviewed, edited, and approved the final version of the manuscript.

FA and DP obtained funding for the studies.

## Acknowledgements

Research was sponsored by the Army Research Laboratory and was accomplished under Cooperative Agreement Number W911NF-21-2-0208 (FA & DP). The views and conclusions contained in this document are those of the authors and should not be interpreted as representing the official policies, either expressed or implied, of the Army Research Office or the U.S. Government. The U.S. Government is authorized to reproduce and distribute reprints for Government purposes notwithstanding any copyright notation herein. The authors would like to thank Drs. Sunil Saxena, David Waldeck, and Michael Hatridge for serving as co-investigators on the funding for this work.

## Data Availability

New data analyses are included in this manuscript and its associated supplement, which will be made available upon publication of this work. The original code is available in Appendix S3 and will be made available online at GitHub once this manuscript is accepted. Since this is a review, the original data input into the models and meta-analyses is available in the original publications as cited in this work.

---

**Figure S1:**
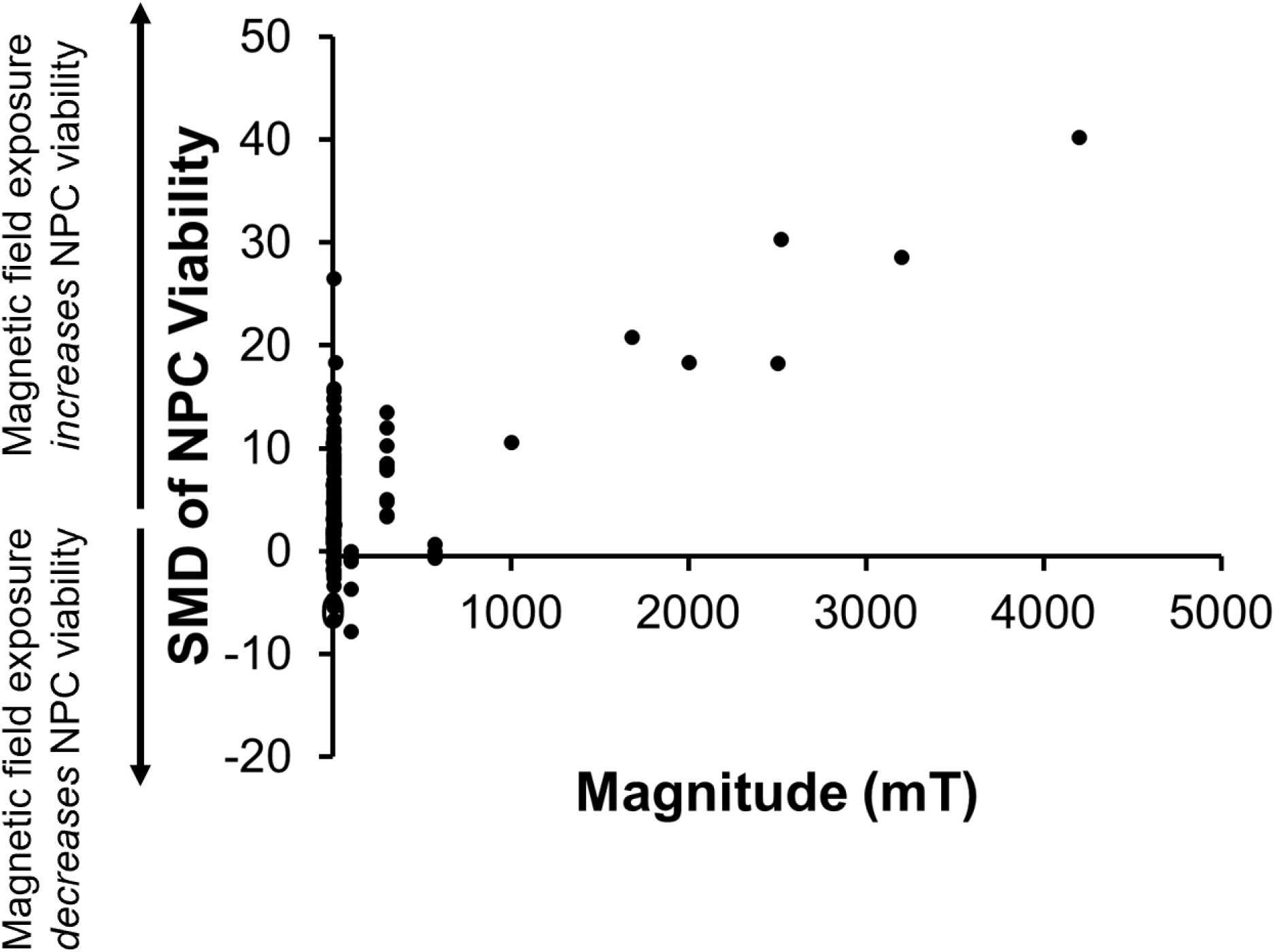
Raw data input into neural stem/progenitor cell (NPC) viability and magnitude model.

